# N6-Adenosine methylation regulates the translation of insulin mRNA

**DOI:** 10.1101/2022.09.20.508712

**Authors:** Daniel Wilinski, Monica Dus

**Affiliations:** Department of Molecular, Cellular, and Developmental Biology, The University of Michigan; Ann Arbor, MI 48109

## Abstract

Relatively little is known about the first step of insulin synthesis: the translation of the mRNA. Here we show that the translation of *D. melanogaster insulin 2* mRNA (*dilp2*) is controlled by methylation of N6-adenosine (m^6^A) in the 3’ UTR. Mutations in the m^6^A writer *Mettl3* and methylated-residues in the *dilp2* 3’UTR decreased the levels of *dilp2* mRNA associated with the polysomes, and the total amount of dilp2 protein produced. This resulted in aberrant energy homeostasis and diabetic-like phenotypes, consistent with the specific function of dilp2 in adult metabolism. Conserved m^6^A signatures were also identified in the 3’ UTRs of vertebrate insulin mRNAs. These data identify m^6^A as a key regulator of insulin protein synthesis and energy homeostasis in metazoans and demonstrate an essential role for m^6^A in translation, with important implications for diabetes and metabolic disease.

**One-Sentence Summary:** The most abundant modification in eukaryotic mRNAs controls the synthesis of insulin protein in *D. melanogaster.*

---

m^6^A is an abundant internal modification of eukaryotic mRNAs involved in regulating mRNA stability, turnover, and translation (*1*). As such, it plays an important role in many biological processes, including differentiation, development, and cancer (*2*). Recent studies have also implicated the m^6^A pathway in energy homeostasis, particularly in the pathogenesis of Type 2 Diabetes (T2D), a chronic disease characterized by the inability to regulate blood glucose via the insulin hormone. Genetic depletion of m^6^A levels in mammalian insulin β-cells resulted in higher circulating glucose and diabetic phenotypes (*3*, *4*) as well as impaired insulin-cell mass and maturation *in vitro* (*4*). Reduced m^6^A levels were also observed in the pancreatic β-cells of humans with T2D (*5*). These observations suggest that m^6^A plays a role in energy homeostasis, especially glucose regulation and T2D. However, the specific mechanisms through which this pathway controls energy balance remain unresolved. Further, translational control is thought to be a critical step in the production of insulin, but the mechanism remains uncharacterized (*6*, *7*). Here we took advantage of the ancestral conservation of the m^6^A (*8*) and insulin systems in invertebrates (*9*) to identify the molecular mechanisms through which m^6^A regulates glucose homeostasis and its contributions to diabetes and metabolic disease.

In *D. melanogaster*, 14 neuroendocrine cells located in the *pars intercelebralis* (Fig. 1A) regulate hemolymph glycemia and energy homeostasis by producing three insulin-like hormones (*10*). While these insulin-like hormones have redundant roles in development, the *Drosophila* insulin-like 2 (dilp2) peptide is necessary and sufficient to regulate hemolymph glycemia and fat levels in adult flies; dilp2 also has the highest homology to human insulin, while other dilps are related to the Relaxin family of hormones (*9*, *11*–*13*). Mutations in *dilp2* and ablation of the insulin-producing cells result in higher levels of circulating sugar and reduced metabolic function, phenotypes that can be rescued by injection of human insulin (*9*–*12*). Networks of transcripts involved in fly metabolism are methylated (*14*, *15*). The accessible location of these cells, together with the genetic and metabolic analysis tools available, makes the fly a uniquely suited model to investigate the *in vivo* function of the m^6^A pathway in glucose and energy homeostasis.

**Fig. 1.**
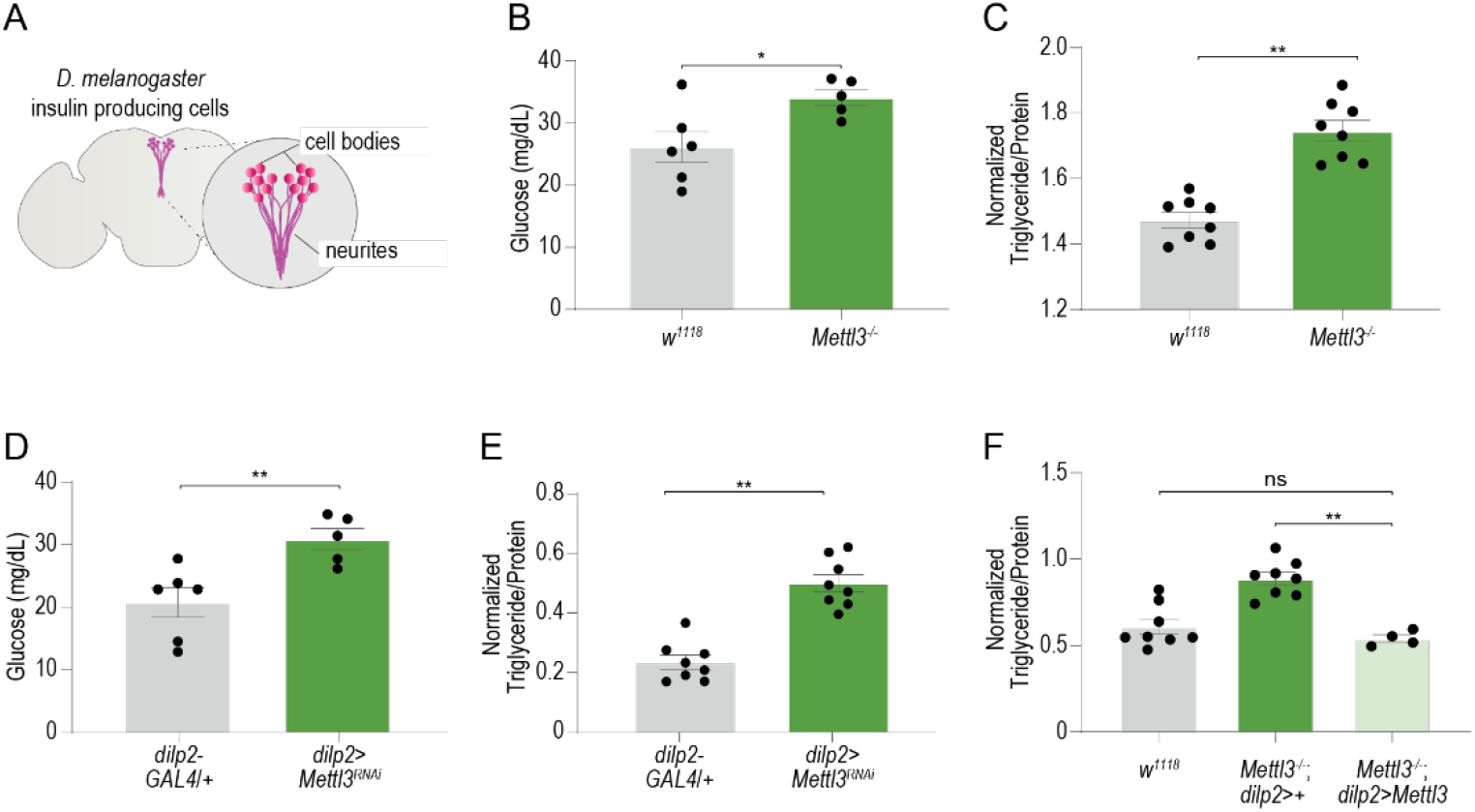
Mettl3 is required for glucose balance and energy homeostasis in the insulin-producing cells. (A) Diagram showing the location and anatomy of insulin-producing cells in the fly brain. (B) The circulating hemolymph glycemia (n=6,) of fasted Mettl3^−/−^ and *w1118* control flies. (C) Triglyceride levels normalized to protein in male w^1118^CS and mutant Mettl3^−/−^ flies. n=8 pools of two flies. (D) The hemolymph glycemia (n=6) of starved *dilp2*>*Mettl3^RNAi^* and transgenic control flies. (E, F) Triglyceride levels normalized to protein in male flies with cell-specific (E) knockdown (*dilp2*>*Mettl3^RNAi^*) or rescue of (F) (*dilp2*>*Mettl3^−/−^*) of Mettl3; n=4-8 pools of two flies. Student’s t-test. Error bars represent standard error of the mean (SEM). *p<0.05, ** p<0.005.

To tackle this question, we first measured the levels of circulating glucose and of body triglycerides in flies mutant for the conserved methyltransferase writer enzyme *Mettl3 (1).* Compared to control flies, *Mettl3^−/−^* mutants showed higher fasted circulating sugar levels (Fig. 1B), as well as increased triglycerides, as previously reported for *dilp2* mutants (*12*) (Fig. 1C). Cell-specific knockdown of *Mettl3* in the insulin-producing cells using the *Dilp2-GAL4* transgene (50% efficiency, Fig S1A) phenocopied the effects of *Mettl3^−/−^* mutants and resulted in higher circulating glucose and triglycerides (Fig. 1D, E). These phenotypes did not arise from developmental alterations in the dilp2+ cells because knocking down *Mettl3* only in post-eclosion, adult *dilp2>Mettl3^RNAi^; tubulin-GAL80^ts^* flies showed the same energy homeostasis phenotypes as *dilp2>Mettl3^RNAi^* animals (Fig. S1B). Importantly, expression of wild-type *Mettl3* only in the insulin cells in an otherwise *Mettl3^−/−^* mutant background rescued the energy balance defects, indicating that the phenotype arises from these cells (Fig. 1F). Thus, *Mettl3* acts in the insulin cells to regulate energy balance; this is similar to the phenotypes of m^6^A writers mutants observed in mice (*3*).

Since the physiological effects of *Mettl3* depletion are reminiscent of those caused by mutations in the *dilp2* gene –but not other insulin-like peptides (*12*)– and murine data implicated potential changes in insulin levels (*3*, *5*), we asked if *Mettl3* affected the amount of this hormone. We found no changes in the abundance of *dilp2* mRNA between *Mettl3^−/−^* and control flies (Fig. 2A); the amount of other insulin-like mRNAs produced in these cells, *dilp3* and *dilp5*, were unchanged (Fig. S2A, B). In contrast, we observed a marked reduction in dilp2 protein in the insulin-producing cells of *Mettl3^−/−^* mutant flies (Fig. 2B, C); no changes in the number and morphology of the insulin cells, or the amounts of the closely related dilp3 hormone were observed (Fig. S2C, D, E). m^6^A regulates both RNA stability/turnover and translation in a context-dependent way (*2*). The observed decrease in dilp2 protein levels without accompanying changes in mRNA abundance suggests that m^6^A may contribute to the translation of the *dilp2* transcript. This may also explain why we do not observe a concomitant transcriptional response from the other dilps (Fig. S2A, B) (*16*). To test the hypothesis that the m^6^A pathway affects the translational status of the *dilp2* mRNA, we fractionated polysomes from control and *Mettl3^−/−^* mutants and measured the amount of *dilp2* transcript associated with each fraction (Fig. 2D, E). In control flies, 89% of the *dilp2* mRNAs cosedimented with polysome fractions, suggesting active and efficient translation (Fig. 2F, gray). Strikingly, this pattern was reversed in *Mettl3^−/−^* flies, where only 19% of *dilp2* mRNA was found in the heavier fractions; instead, 80% of this mRNA was associated with early fractions, representing ribosomal individual subunits and monosomes (Fig. 2F, green). This suggests that m^6^A is required for loading the *dilp2* mRNA onto polysomes and therefore proper translation of the *dilp2* mRNA into protein.

**Fig. 2.**
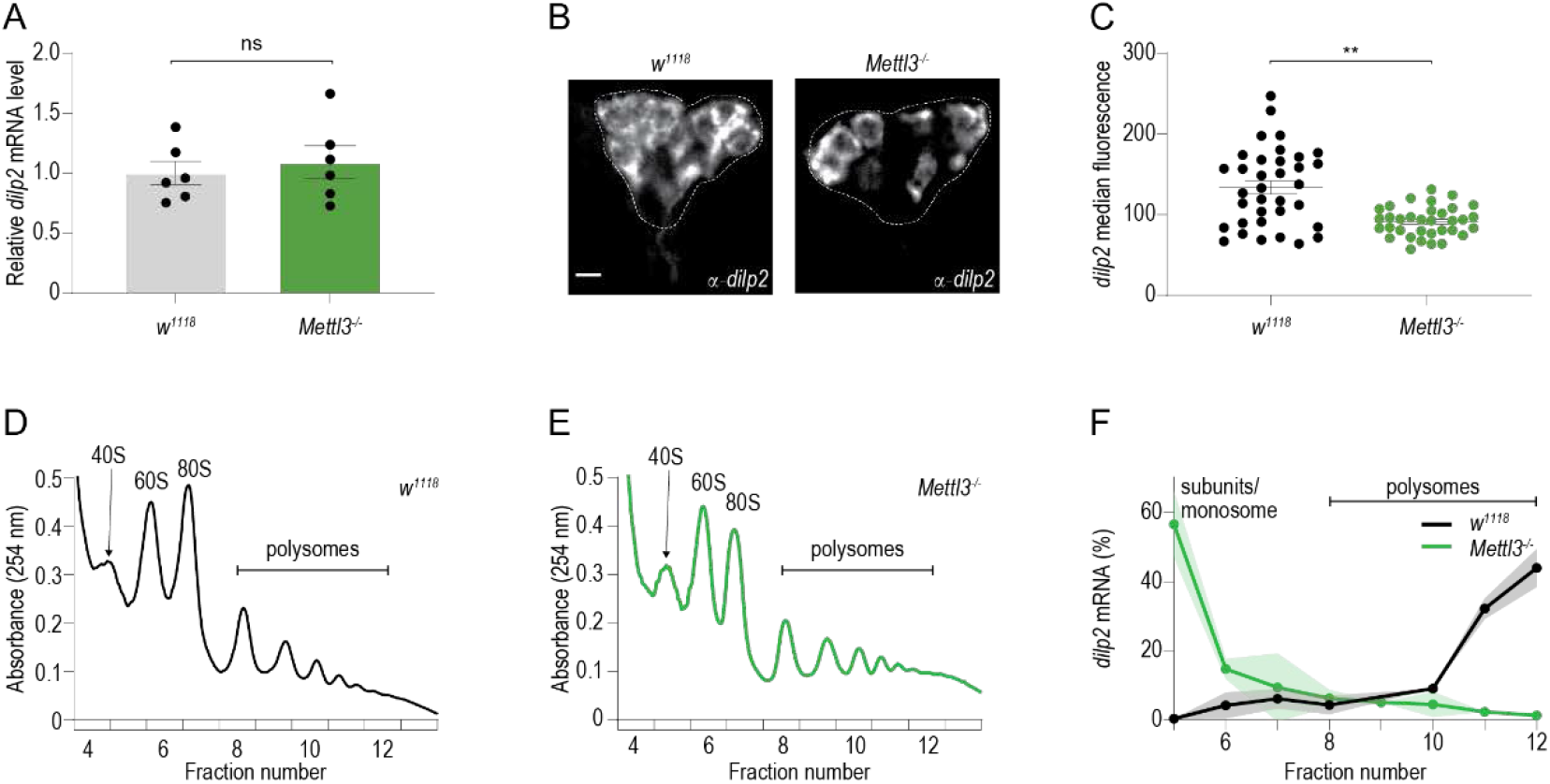
Mettl3 is required for the translation of *dilp2* mRNA. (A) Quantification of *dilp2* mRNA from control and *Mettl3*^−/−^ mutant flies. n=6 sets of 20 flies. Error bars SEM in this and all subsequent panels. Student’s t-test. ns =not significant (p=0.5944). (C) Representative confocal images of immunofluorescence of *dilp2* protein in control (w^1118^CS) and *Mettl3*^−/−^ mutant flies. Scale bar, 20um. (D) Quantification of median dilp2 fluorescence (arbitrary units) of individual insulin-producing cells from *Mettl3* mutants and control flies; n=15 brains. Male flies were starved for 16 hr prior to dissection and collected in 3 independent sets. Student’s t-test. ** p < 0.005. (E-F) Representative polysome profile from sucrose gradient of (E) control (w^1118^CS) and (F) *Mettl3*^−/−^mutant fly heads (400). (G) The proportion of *dilp2* mRNA in sucrose gradient fractions 5-12 normalized to spike-in RNA from control (*w^1118^CS*) and *Mettl3*^−/−^ mutant flies. n=2, 400 heads per sample.

To characterize the molecular causes of this phenotype, we used fly heads to perform m^6^A ultraviolet light-induced Cross-linking and Immunoprecipitation (miCLIP (*17*)), a technique that identifies transcripts marked by m^6^A. The three biological replicates showed good reproducibility with a mean correlation of 0.95 (Fig. S3A) and identified 4,506 m^6^A peaks corresponding to 1,828 genes involved in brain processes including development and plasticity (Data S1 and Fig. S3B). Forty-four percent (2009) of our peaks overlap with previous CLIP data from fly heads including *Atpα* (*18*) (Fig. S3C). Metagene analysis revealed that most m^6^A peaks mark the 5’ <u>u</u>n<u>t</u>ranslated <u>r</u>egion (UTR) of transcripts, while a much smaller portion is present in the 3’ UTR, particularly near the stop codon (Fig. 3A); this is consistent with previous miCLIP studies in flies (*18*, *19*). We also observed an enriched fly RRAC (R=purine) sequence motif at C-to-T crosslinking-induced mutation sites (CIMS) which represent m^6^A sites from CLIP/RIP data (*14*, *17*–*19*) (Fig. 3B, Data S1).

**Fig. 3.**
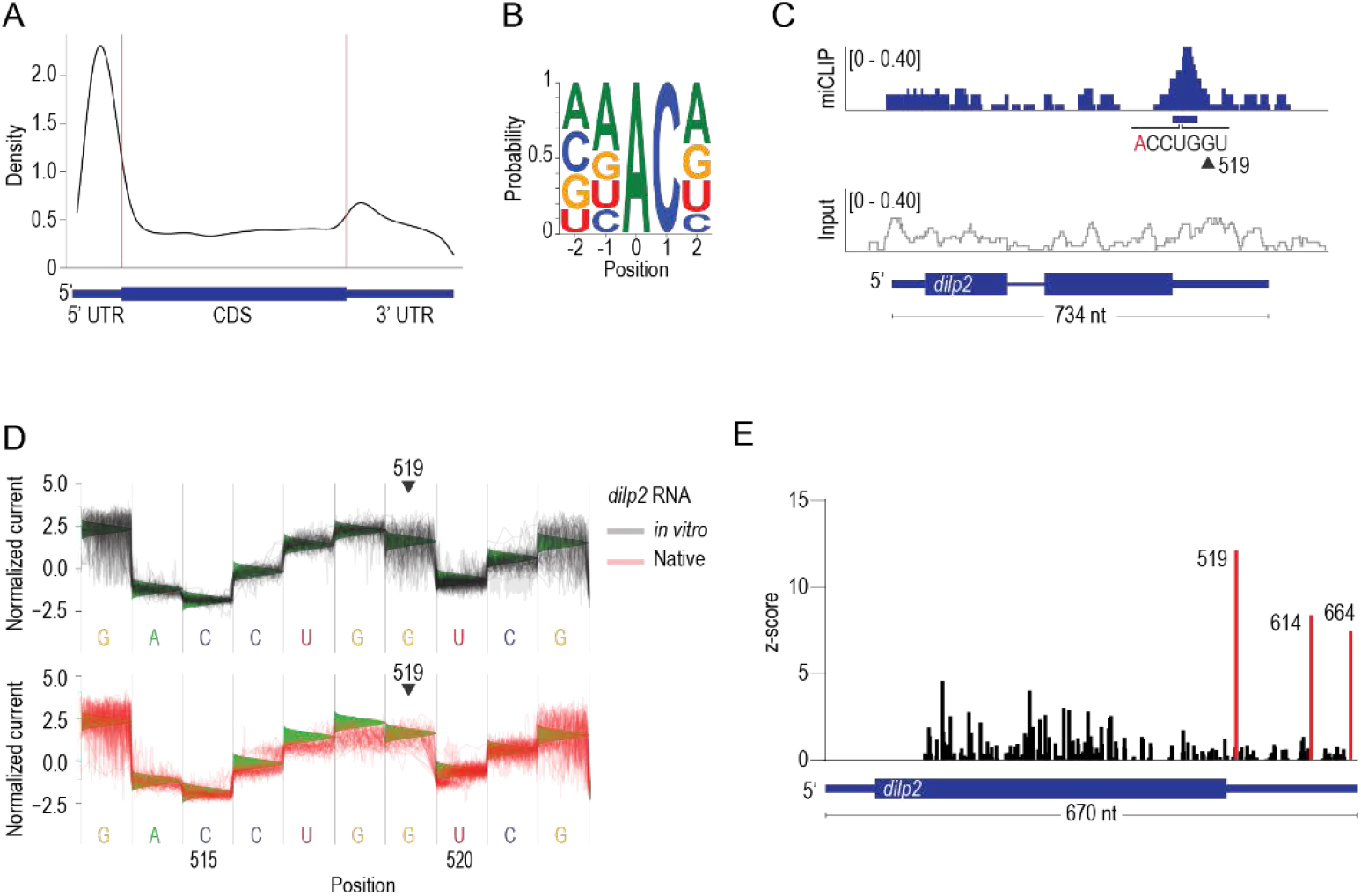
The 3’ UTR of the *Drosophila melanogaster dilp2* mRNA is methylated. (A) Metagene plot of CLIP peaks from *D. melanogaster* head mRNA. Representing the position of 4,506 CLIP peaks. (B) The sequence context of 5,485 CIMS contained within CLIP peaks. (C) CLIP (top, blue) and the no IP input control (bottom, grey) traces mapped to the dilp2 locus (<u>R</u>eads <u>P</u>er <u>M</u>illion mapped reads, RPM). The blue horizontal bar indicates the CLIP peak (FDR <0.05) and the base composition flanking a putative m^6^A site near position 519. (D) Normalized direct-RNA sequencing signal derived from *in vitro* transcribed *dilp2* RNA (top, gray, n=50 reads) and native *dilp2* mRNA from fly heads (bottom, red n = 50 reads) of the region corresponding to the CLIP peak in (C). Green triangles represent the expected current level based on the base calling model (see methods). (E) EpiNano significance trace showing the nucleotide positions that are significantly different (red bar, z-score > 6) between native and *in vitro* transcribed *dilp2* RNAs.

Since we observed lower translation of the *dilp2* mRNA in *Mettl3^−/−^* mutants, we asked if this transcript was methylated. Indeed, an m^6^A peak was present in the 3’ UTR of *dilp2*, shortly after the stop codon (Fig. 3C); none of the mRNAs encoding for other insulin-like peptides expressed in the insulin-producing cells, such as *dilp3* or *dilp5*, showed any signatures of methylation (Fig S3D, E). Thus, the *dilp2* mRNA is methylated *in vivo*. To better characterize the location of the modified A we turned to direct-RNA sequencing (Oxford Nanopore) where modified bases can be detected as changes in normalized current through the nanopore (*20*–*22*). We first *in vitro* transcribed RNA oligomers containing only one methylated or unmethylated A, sequenced them on the Nanopore, and then used the EpiNano algorithm to assess deviations in base-calling between these two identical oligomers (Fig. S4A, Table S1) (*21*). Direct-RNA sequencing of these RNAs revealed a shift in the raw current near the methylated bases (Fig. S4B) which results in a significant difference in base-call “errors” (Fig. S4C). Thus, we can detect N6-methylation as a deviation in base calling at or near to the methylated A via direct-RNA sequencing. To define if the native *dilp2* mRNA was methylated *in vivo*, we enriched for insulin-cell-specific mRNAs by expressing the FLAG-tagged Ribosomal Protein 3 *UAS-<u>RPL</u>3::FLAG* transgene with *Dilp2-GAL4*. We then direct-RNA sequenced the purified poly(A)+ RNA and defined deviations between base calls of *in vitro* transcribed and native RNA (*21*). This experiment revealed a significant difference in the 3’ UTR between the *in vitro* transcribed and native *dilp2* RNAs (Fig. 3D, E). In particular, three bases in the 3’UTR (Positions G519, C614, C664) had significantly higher base-call deviations in the native RNA compared to the *in vitro* transcribed *dilp2* RNA. Position G519 corresponds to the CLIP peak and positions C614 and C664 are part of AC dinucleotides (Table S2). Together with the results from miCLIP, these data show that at least three specific ACs in the 3’UTR of the *dilp2* mRNA are methylated *in vivo* in the *D. melanogaster* insulin-producing cells.

To investigate whether N-6 RNA methylation of the *dilp2* 3’ UTR directly affects its translation, we generated flies that lacked “methylateable” A’s in the 3’ UTR (*dilp2* ^m6A−/−^). Using the miCLIP and Nanopore analyses, we selected 11 “AC” nucleotides and mutated these into “UC” using CRISPR (Fig. S5A). These included the AC near G519 and C614 with the strongest methylation signals (Fig. 3C, E); A663 could not be removed for technical reasons. Clone 364-4 was selected for further study after confirming all the A>U mutations by Sanger sequencing (Fig. S5B). Sucrose density polysome gradients showed that compared to control flies, where 92% of the *dilp2* mRNAs were in heavier fractions, representing the polysomes, 69% of *dilp2* mRNAs from *dilp2* ^m6A−/−^ flies were associated with early fractions (Fig 4A, S6A, B). Consistent with this, quantification of the total levels of dilp2 protein with anti-dilp2 specific antibodies in the insulin cells revealed a 20 % decrease in *dilp2* ^m6A−/−^ flies compared to controls (Fig. 4B, C). Strikingly, we found that *dilp2* ^m6A−/−^ mutants recapitulated the deficits in glucose and energy homeostasis observed in *Mettl3* mutant flies, with an increase in fasting glucose levels and in triglycerides (Fig 4 D, E). Together, these findings suggest that, in flies, m^6^A modification of the *dilp2* mRNA controls the effective translation, and thus synthesis, of the insulin protein.

**Fig. 4.**
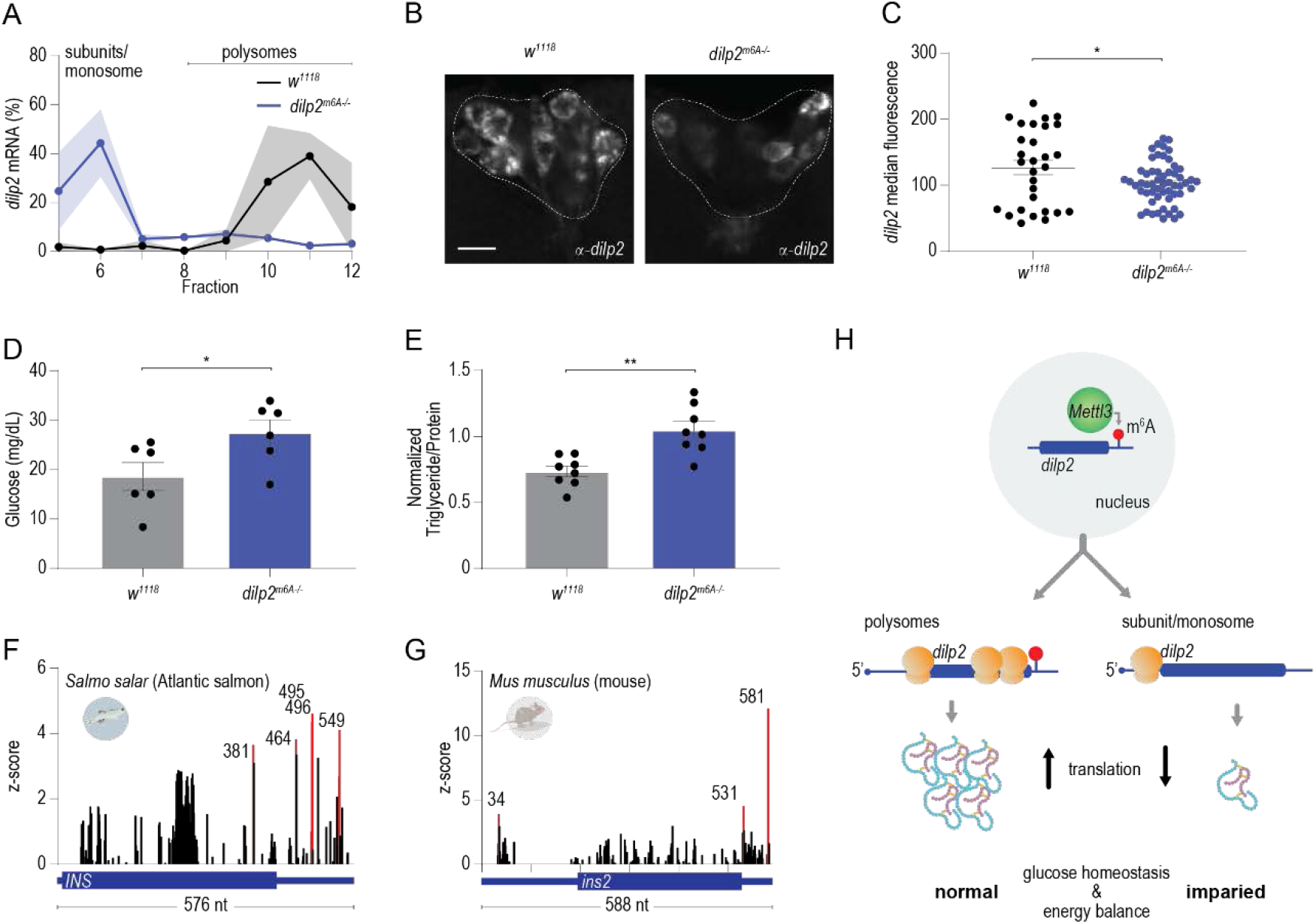
Methylation of dilp2 mRNA is necessary for robust translation. (A) Proportion of *dilp2* mRNA found in polysome gradient fractions 5-12 compared to a spike-in RNA. n=3 samples of 400 heads each. Shading represents SEM. (B) Representative confocal images of dilp2 protein in fasted control and *dilp2* ^m6A−/−^ mutant flies. (C) Quantification of median *dilp2* fluorescence of individual insulin-producing cells from 6 brains from B). Scale bar, 20um. Error bars SEM. * p<0.01 Student’s t-test. (D) The circulating hemolymph glycemia (n=6) of fasted *dilp2* ^m6A−/−^ and control flies. (E) Triglyceride levels normalized to protein in control (w^1118^) and mutant *dilp2^m6A−/−^* flies. n=8 pools of two flies. (F, G) Direct RNA sequencing significance trace comparing *in vitro* transcribed salmon INS (F) or mouse INS2 (G) RNA to native mRNA isolated from salmon pancreatic tissue (F) or mouse pancreatic islets (G). Numbers represent the significantly different nucleotides (red bar, z-scores > 4 or >5) between native and *in vitro* transcribed RNAs. (H) Model of translational control of *dilp2* RNA via m^6^A. *p < 0.05, ** p < 0.005.

Elements in the 3’ UTR of mammalian insulin are thought to play an important role in its translation (*6*, *7*). Given the conservation of the insulin and m^6^A pathways in metazoa, we asked if signatures of m^6^A were also present in vertebrate insulin mRNAs. To do this, we obtained polyA-selected mRNAs from Atlantic salmon pancreatic tissue (*Salmo salar*) and mouse islet cells and analyzed them by direct-RNA sequencing. We analyzed 5,000 and 10,000 reads that mapped to the INS and INS2 insulin genes from salmon and mouse, respectively. This analysis revealed one putative m^6^A site in the 3’UTR of salmon insulin (Fig. 4F, Table S2, C464) and two putative m^6^A sites in INS2 3’ UTR (Fig. 4G, Table S2, A531, and C581). This suggests that regulation of insulin translation by m^6^A may be conserved.

Taken together, our observations support a general model where methylation of adenosines in the 3’UTR of the *dilp2* mRNA enhances translation by promoting the association of this mRNA with polysomes which is consistent with a possible defect in translation initiation (*23*). In contrast, m^6^A in the 5’ UTR has been linked to non-canonical translational control (*24*). Although we cannot exclude that this mechanism also controls other aspects of *dilp2* mRNA, our polysome profiling data links m^6^A marks in the *dilp2* mRNA with translational initiation. In the absence of this epitranscriptomic signal, less insulin is produced, compromising the animal's ability to regulate sugar and energy homeostasis. Interestingly, this mark only controls the translation of the *dilp2* mRNA and not the closely related *dilp3* and *dilp5* genes, consistent with the fact that the deposition of the signal is specific (*25*). Further, it suggests that previously observed transcriptional compensations among different insulin-like peptides in *D. melanogaster* are tied to mRNA abundance, not protein levels (*16*).

Based on our discovery that adenosines in the 3’UTR of mouse and salmon insulin mRNAs are marked with m^6^A, and on previous data showing that Mettl3 and Mettl14 are involved in glucose homeostasis, we anticipate that regulation of insulin translation and energy homeostasis by this RNA modification will be a conserved mechanism in vertebrates. Since m^6^A has been linked to cellular metabolism and signaling, this mode of regulation could provide a mechanism to time insulin translation with the presence of specific physiological signals, as well as to prepare for its fast production. This is consistent with findings that elements in the 3’UTR of mammalian insulin are important for its regulated production (*6*, *7*) and that altered levels of m^6^A are found in the islets of people with T2D (*5*). However, it is still unclear if the m^6^A mark is dynamic and could really be used to switch translation. However, rather than toggling translation on and off, m^6^A could promote protein synthesis by direct cellular compartmentalization via phase separation (*1*). This work also provides direct evidence for the hypothesis that m^6^A in the 3’ UTR plays an essential and robust role in translation, although the mechanisms remain to be defined (*1*). Besides its relevance to the study of insulin and physiology, the methylation of *dilp2* mRNA could be a viable model to address these and future questions about the biology of m^6^A. In summary, our work has uncovered a new fundamental step in the biosynthesis of insulin and the biology of m^6^A, with critical implications for understanding RNA control (or translational control) mechanisms in the regulation of energy homeostasis and the etiology of metabolic disease.

## Acknowledgments

**General**: We thank Pierre Léopold for the kind gift of Dilp2 antibody, Patrick Callaerts for the gift of Dilp3 antibody, Randy Seeley for mice and Corentin Cras-Méneur for isolating Mouse islets, Jean-Yves Roignant for *Mettl3* mutant flies, and the Blooming Drosophila Stock center for other flies used in this study. Salmon tissue was a gift from Brian Peterson (USDA); we thank Cunmin Duan for advice. We are grateful to Peter Todd and Shannon Miller for training and for the use of their polysome fractionation equipment and training; Christopher Lapointe for thoughtful comments on the manuscript. We also thank Julia Kuhl for designing some of the graphics in this manuscript.

## Funding

NIH R00 DK-97141 (MD)

NIH 1DP2DK-113750 (MD)

Rita Allen Foundation (MD)

NSF CAREER 1941822 (MD)

NIH T32 DA007268 (DW)

NIH P30 DK089503 (MD and DW)

## Author contributions

Conceptualization: DW and MD

Methodology: DW

Investigation: DW and MD

Visualization: DW and MD

Funding acquisition: MD

Project administration: MD

Supervision: MD

Writing – DW and MD

Writing – review & editing: DW and MD

## Data and materials availability

All data are available in the main text or supplementary materials. RNA-sequencing reads were deposited in Gene Expression Omnibus accession number GSE207547. (The following secure token has been created to allow review of record GSE207547 while it remains in private status: cfkreeumnfajzkn) *dIlp2* 3’UTR mutants are available upon request.

## Supplementary Materials

### Materials and Methods

#### Fly lines and husbandry

All flies were maintained at 25 °C in a humidity-controlled incubator with a 12:12 hr light/dark cycle. Animals were fed Bloomington Food B (cornmeal-glucose) ad-lib and provided fresh food every other day. For all experiments, flies were collected under CO2 anesthesia, 2–4 days following eclosion, and housed in groups of 20–30 to age until testing (6-10 days old). The stocks used are listed in Table S3. Depending on the genetic background of the mutations, we used *w^1118^* (Rainbow Transgenic Flies, Camarillo, CA) or *w^1118^ Canton-S* flies (Benzer lab, Caltech) as controls.

#### RNA extraction

Standard methods to isolate total RNA were used for quantitative polymerase chain reaction (qPCR), crosslinking and immunoprecipitation (CLIP), and direct-RNA sequencing. Briefly, heads from 10 to 20 flies were dissected and immediately frozen on dry ice for qPCR. 800 fly heads were isolated from bodies by sieving for CLIP and direct RNA-sequencing. All samples were stored at −80 °C until extraction. Phenol-chloroform (Invitrogen, 15596018) and RIPA buffer (150 mm NaCl, 1% Nonidet P-40, 0.5% Sodium deoxycholate, 0.1% SDS, 50mM Tris pH 7.5) were added to frozen samples and homogenized by bead bashing (Bead Ruptor, 19-040E). Phenol-chloroform extracted RNA was precipitated by isopropanol with GlycoBlue coprecipitant (Invitrogen AM9515). RNA was stored at −80 °C until further processing.

### CLIP

#### Library preparation

We adapted miCLIP (*17*) to use infrared-dye-conjugated irCLIP adaptors (*26*). Briefly, 20 ug of poly(A) selected (Invitrogen, 61012) RNA for each biological replicate from 800 heads of *w^1118^CS* flies was used as input to the CLIP, no antibody control, and input only (no CLIP) reactions. Fragmented RNA (Ambion, E6150S) was incubated for 2 hr at 4 °C with antibody against m^6^A (Abcam, ab151230), UV-crosslinked, then immunoprecipitated with magnetic beads (Invitrogen, 10004D) as in (*17*). Pre-adenylated dye-conjugated linkers were ligated (NEB, M0351S) to dephosphorylated 3’ ends of RNA fragments overnight at 16 °C (Table S1). RNA was extracted with NuPAGE SDS buffer and separated by NuPAGE gel (Invitrogen, NP0321), transferred to nitrocellulose membrane and visualized for excision at 800 nm. RNA-antibody complexes were released from the membrane by proteinase K digestion and RNA was purified with phenol-chloroform extraction. RNA was reverse transcribed (Invitrogen, 18080085), circularized (Lucigen, CL4111K), and PCR-amplified (Thermo Scientific, F530S) following the published protocol (*17*). Libraries were subjected to 151 bp paired-end sequencing according to the manufacturer’s protocol on an Illumina NovaSeq 6000 at the University of Michigan Genomics core.

#### Bioinformatics

Sequencing reads were de-multiplexed using the Bcl2fastq2 Conversion Software (Illumina). Five prime end unique molecular identifiers (UMI, 9 nt random sequence) were used to remove PCR duplicates with a custom script. Then UMI and sequences and sequencing adapters were removed (fastx_clipper, 0.0.14). Reads were mapped to the Ensembl *D. melanogaster* genome (BDGP6) genome using Star (v 2.7.5a, (*27*)) with default settings (Table S4). Aligned reads were peak-called using Piranha (v 1.2.1) (*28*) (Data S1). Metagene analysis was performed using MetaPlotR (*29*). Crosslink induced mutation site analysis we performed as in (*17*). Sequence logo was generated using weblogo (v 2.8.2, (*30*)). Gene ontology (GO) analysis was performed using Gene Set Enrichment Analysis (GESA) implemented in the ClusterProfiler R package version 4.2.2 (*31*). We used Benjamini–Hochberg to correct for multiple hypothesis testing and “biological process” from the Bioconductor package “Genome Wide Annotations for Fly” version 3.8.2 for GESA (*32*).

### Polysome fractionation

Polysome profiles were performed as previously described (*33*). Briefly, 300 fly heads were homogenized using a bead beater (Bead Ruptor, 19-040E) in 800 ul polysome extraction buffer (300 mM NaCl, 50 mM Tris-HCL (pH 7.5), 10 mM MgCl2, 200 mg heparin/ml, 400 U RNasin/ml, 1.0 mM, phenylmethylsulfonyl fluoride, 0.2 mg/ml cycloheximide, 1% Triton X-100, 0.1% sodium deoxycholate) then incubated on ice for 10 minutes. Lysate was cleared by centrifugation at 10,000 x g for 10 min at 4 °C. Equal A260 units were layered onto a 10–50% sucrose gradient in resolving buffer (20mM Tris-HCl (pH7.5), 150mM NaCl, 15mM MgCl2, 1mM DTT, 100ug/mL cycloheximide) and separated using a Beckman SW41Ti rotor (30,000 rpm for 3 hr at 4 °C). The absorbance (254 nm) was monitored and 750 ul fractions were collected using a Brandel pump set to a flow rate of 1.5mL/min. Equal molar concentrations of *Saccharomyces cerevisiae* enolase-2 (*Eno2*) transcript was added to all fractions before RNA isolation. Nucleic acid was precipitated from each fraction then pellets were resuspended in water and phenol chloroform extracted following the manufacturer's protocol (Invitrogen, 15596026). The RNA was precipitated with isopropanol and GlycoBlue. RNA was resuspended in 10 ul water and equal volumes from each fraction were reverse transcribed following the manufacture recommendations (Invitrogen, 18080085). Fractions 5-12 were probed for *dilp2* and *Eno2* by qPCR (Table S1).

### qPCR

Reverse transcription was performed using SuperScript III (Invitrogen, 18080085) with 1 ug of total RNA as input and primed with oligo(dT) (Invitrogen, 18418012) according to the manufacturer's protocol for transcript abundance analysis. Quantitative-PCR was performed following manufacturer's directions (Applied Biosystems, 4367659) for all experiments. Primers were added at a 2.5 μM concentration in 20 ul reactions. Reactions were run on the StepOnePlus Real-Time PCR System (Applied Biosystems), and quantifications were normalized relative to the reference gene ribosomal protein 49 (*Rp49*) for transcript abundance or spike-in, *Eno2*, for polysome fractionation (delta Ct method).

### Fat and lean mass analysis

Colormetric measurements of triglycerides (Stanbio, SB-2100-430) and protein (Pierce, PI23225) were done as previously described (*34*), where 1 biological replicate n=2 male flies. Flies were collected 3-5 days post eclosion and homogenized with a Bead Ruptor. Standard curves were generated for each to normalize the concentration of samples. Samples were quantified using a Tecan Spark plate reader at 562 nm for protein or 500 nm for triglycerides.

### Immunofluorescence

Immunofluorescence was done essentially as (*35*). Male flies 3-5 days post eclosion were sorted then aged for 2-5 days on standard food. Flies were then fasted in vials with a wetted Kimwipe for 16-18 hours before dissection. Brains were dissected in PBS, fixed (4% paraformaldehyde aqueous solution in 1X PBS), blocked (10% normal goat serum, 2% Triton X-100, 1X PBS), and then incubated overnight in primary antibody (rat anti-dilp2 (1:500) (*36*), rabbit anti-dilp3 (1:500) (*37*)). After washing (3% NaCl, 1% Triton X-100, 1X PBS), brains were then incubated at room temp overnight in secondary antibody: either goat anti-rat Alexa Fluor 647 (Invitrogen, A-21247) or goat anti-rabbit Alexa Fluor 488 (Invitrogen, A-11008). Brains were mounted in FocusClear (CelExplorer, FC-101) on coverslips, and the cell bodies were imaged using an FV1200 Olympus confocal with a 20x objective. Median intensity of individual insulin-producing cells was quantified using Fiji (*38*).

### *in vitro* transcription for direct-RNA sequencing

DNA insulin templates were synthesized by IDT based on transcripts FBtr0076329 (fly *Dilp2*), NM_001185083.2 (mouse Ins2), and two randomized sequences containing only one adenine (Supplemental Table 2). To generate the salmon Ins template, we PCR amplified (Thermo Scientific, F530S) DNA from cDNA and Sanger sequence-verified the product (XM_014198195.2). Each template DNA sequence was used for *in vitro* transcription (Invitrogen, AM1334). To generate the randomized DNA template “rand-A” the reaction mixture contained bases adenosine, uracil, cytosine, and guanosine while the randomized DNA template “rand-m6A” used N6-methyladenosine in the palace of adenosine. All the other *in vitro* transcription reactions used only the standard bases. The reactions were performed following the manufacturer's protocol overnight.

### Direct-RNA sequencing

500 ng of poly(A) selected (Invitrogen) RNA from heads of flies or a total 500 ng of pooled *in vitro* transcribed RNA was used for library preparation following the manufacturer's protocol (Oxford Nanopore, SQK-RNA002, Version DRS_9080_v2_revB_22Nov2018). Briefly, the RT adapter was ligated to the RNA, reverse transcription was performed, and the RNA-cDNA hybrids were purified. Next, the second adapter was ligated to the RNA and again purified. The libraries were loaded onto the MinION flowcell (R9.4.1). The Oxford Nanopore sequencer was run for 24-36 hours. Data were base called using Oxford Nanopore’s Guppy (v 3.1.5) and aligned to the reference sequences using MiniMap2 -ax splice -uf -k14 (v 2.17) (*39*). Only reads that passed filtering and that mapped to the reference were considered for further analysis. Next, aligned reads from biological samples (modified) and matched *in vitro* transcribed RNAs (unmodified) were used as input to EpiNano (EpiNano-Error, v 1.2) to determine the positions of modifications (*21*). We plotted the data from the longest transcript (NM_001185083.2) for EpiNano mouse data. The Tombo suite of tools was used to visualize reads (*40*).

### Circulating glucose assay

Hemolymph was collected from starved (12-16 hours) male flies from 40-50 flies per replicate by centrifugation. Circulating glucose levels were measured as previously described (*41*). Briefly, 0.5 ul of hemolymph was added to 100 ul of HexoKinase (HK) reagent (Sigma, GAHK20), incubated at room temperature for 15 min, then absorbance at 340 nm was measured on a Tecan Spark plate reader.

### Generation of *dilp2* m^6^A mutant flies

CRISPR constructs were synthesized and micro-injected into *w1118* flies by Rainbow Transgenic Flies (Camarillo, CA). The pScarless donor vector (dsRed marker) was introduced to remove the endogenous Dilp2 3’UTR and replace it with a mutant Dilp2 3’UTR that replaced 11 AC dinucleotides to TC (Supplement Fig. 4A). F1 progeny were screened for transformation with dsRed fluorescence. Positive transformants were Sanger sequenced (below) to verify the correct insertion.

### Sanger Sequencing

Genomic DNA was extracted from two male flies from each fly line with positive dsRed expression by silica column purification (Invitrogen, K182002). The *dilp2* locus was PCR amplified (Thermo Scientific, F-530XL), PCR products were purified and normalized to 5ng/ul with 10 pmol/ul of the with appropriate primer added. Samples were submitted to Eurofins Genomics for sequencing and traces were analyzed with Benchling software.

### Mouse islet isolations

Islet cells were collected from 4 fasted male mice from Vil-Cre backcrossed to C57 9x by the University of Michigan Islet Isolation core. The pooled tissue was added directly to Trizol (Invitrogen, 15596018) and stored at −80 °C.

### Salmon pancreatic tissue isolation

The Atlantic salmon used was approximately 2 years old and post-smolt. The pancreatic tissue from one individual was isolated from the surrounding pyloric caeca. Upon removal, the tissue was immediately added to Trizol and stored at −80 °C.

**Supplementary Fig. 1.**
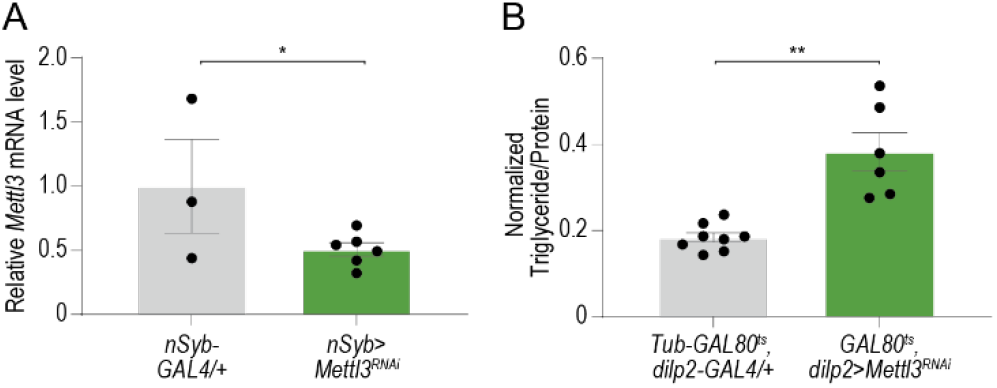
The effects of *Mettl3* KD on energy homeostasis are not developmental. (A) Quantification of *Mettl3* mRNA from heads of control *Dilp2>w1118cs* and *Dilp2*>*Mettl3^RNAi^* flies. n=3, 5 sets of 20 flies. (C) Triglyceride levels normalized to protein in male control (*Tubulin-GAL80^ts^, dilp2-GAL4 / +*) and *Tubulin-GAL80^ts^; dilp2>Mettl3^RNAi^* flies. n=8, 6 pools of two flies. Error bars are SEM. Student’s t-test. * p < 0.05

**Supplementary Fig. 2.**
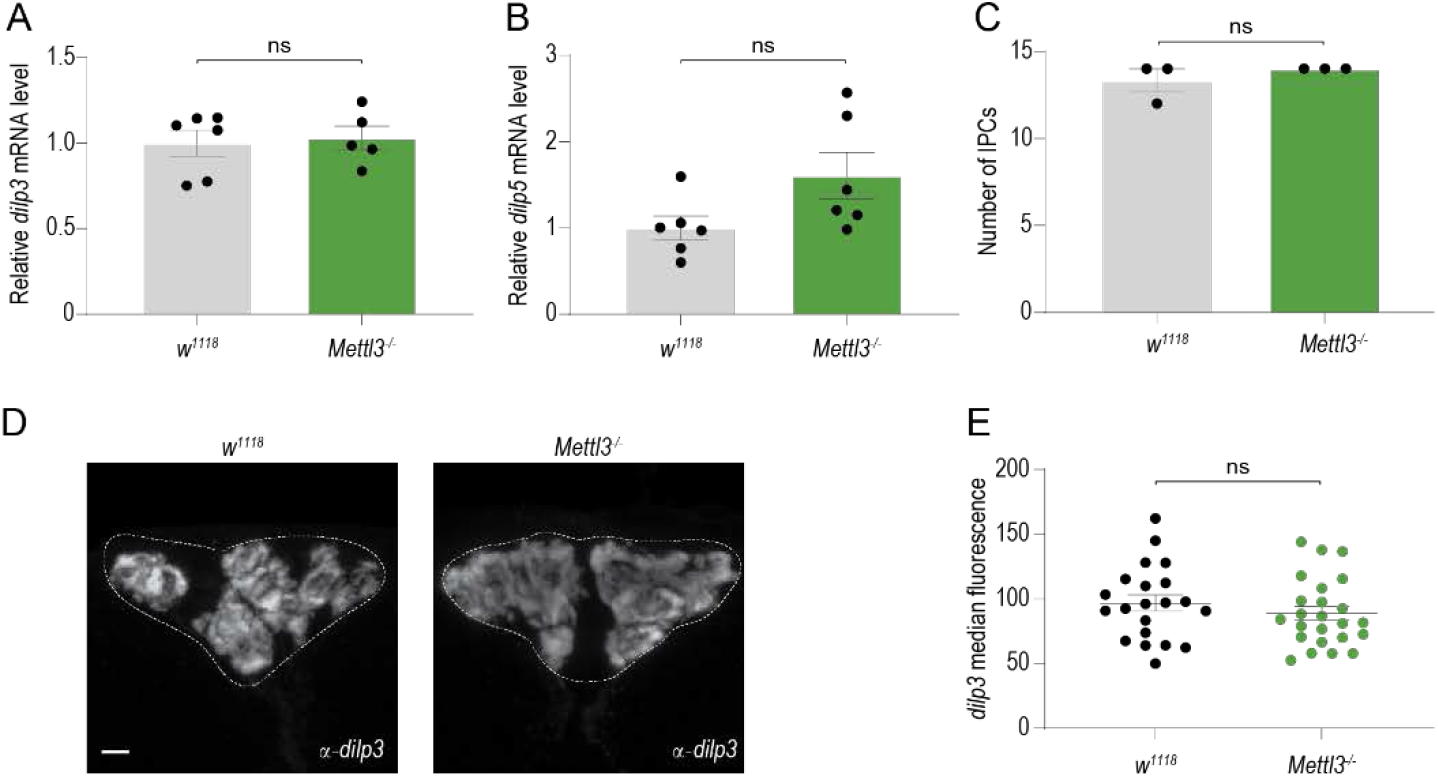
Additional phenotyping of *Mettl3* mutants. (A, B) Quantification of (A) *dilp3* and (B) *dilp5* mRNA from heads of control (*w^1118^CS*) and *Mettl3^−/−^* mutant flies. n=6 sets of 20 flies. (C) Representative confocal images of immunofluorescence of *dilp3* protein in control (*w^1118^CS*) and *Mettl3*^−/−^ mutant flies. Scale bar, 20um. (D) Quantification of median dilp3 fluorescence of individual insulin-producing cells from C), n=6 per genotype. Scale bar, 20um. (E) Error bars are SEM. Student’s t-test. ns = not significant, * p < 0.05.

**Supplementary Fig. 3.**
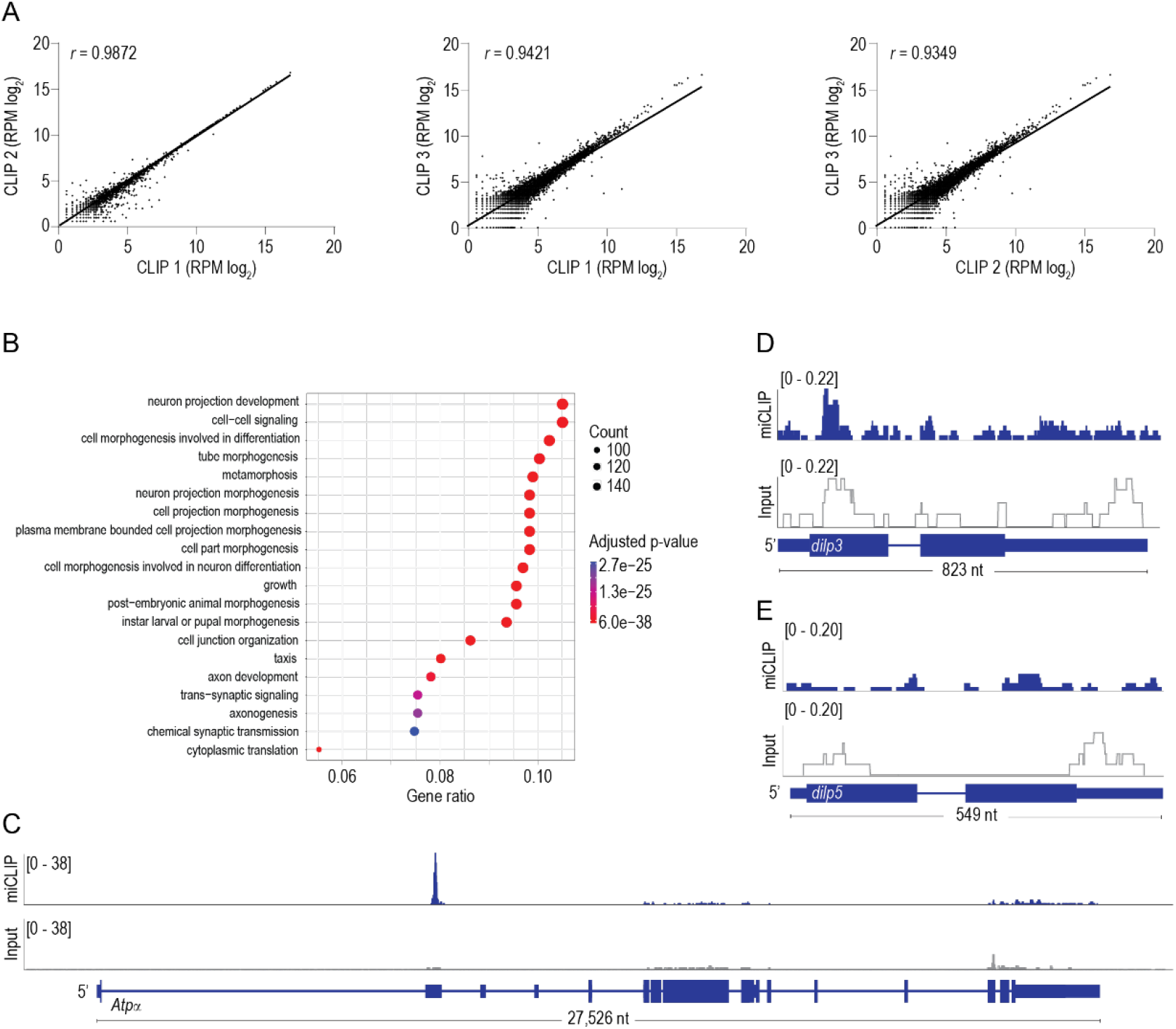
Reproducibility of biological CLIP replicates. (A) Correlation plots of log2 normalized reads per CLIP peak. Each dot represents a CLIP peak found in all three biological replicates. Pearson’s correlation coefficient (*r*). (B) Gene ontology (GO) enrichment analysis of genes that harbor a CLIP peak. Circle size represents the number of genes that have CLIP peaks in the corresponding GO categories. The color represents the significance of the enrichment (Benjamini–Hochberg corrected p-value from Gene Set Enrichment Analysis (GSEA)). (C, D, E) CLIP (blue) and input (gray) traces mapped to the *Atpα* (C), *dilp3* (D) and *dilp5* (E) loci (RPM).

**Supplementary Fig. 4.**
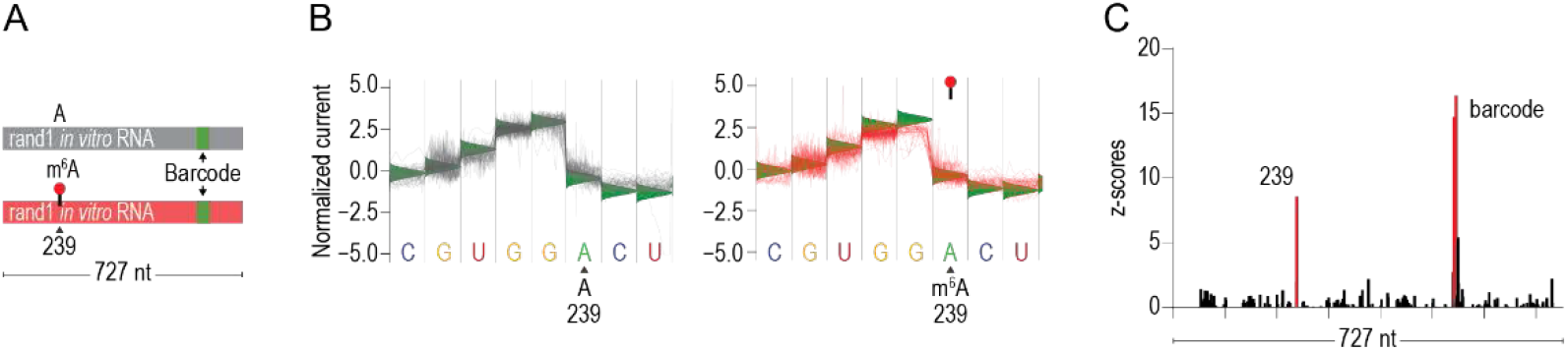
Direct RNA sequencing of *in vitro* transcribed control RNAs. (A) Schematic of the randomly generated random-1 (rand1) *in vitro* transcribed RNAs. The RNAs were transcribed with A (gray) and with m^6^A (red). The sequences were identical except for the 6 nt molecular barcode depicted by the green block to unambiguously distinguish between the unmethylated and methylated RNA. (B) Normalized direct RNA sequencing signal derived from direct RNA sequencing of *in vitro* transcribed RNA with A (gray) and with m^6^A (red). n=50 reads plotted. Green triangles represent the expected current level based on the base calling mode used by Guppy (Version 3.1.5, Oxford Nanopore Technologies). (C) EpiNano significance trace across the *in vitro* transcribed RNA sequence. Significant position 239 (red) corresponds to the base following the methylated A (238). Other significant bases labeled “barcode” correspond to the green barcode in (A).

**Supplementary Fig. 5.**
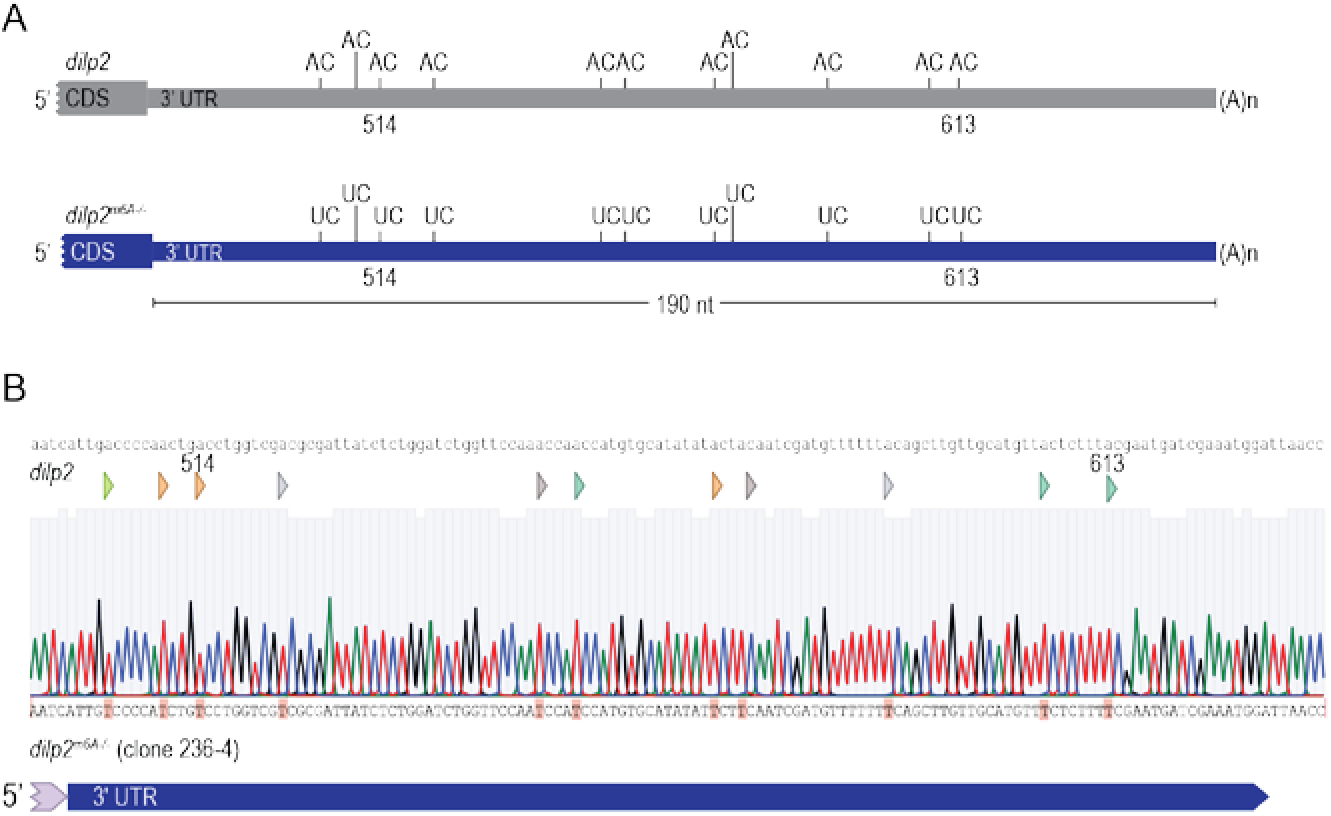
Creation of *dilp2^m6A^* mutant flies. (A) Diagram of CRISPR strategy to replace 11 AC dinucleotides in 3’ UTR of the *dilp2* transcript (dilp2^m6A−/−^gray). (B) Sanger sequencing of genomic DNA from positive clone 364-4 showing all 11 AC dinucleotides replaced with UC.

**Supplementary Fig. 6.**
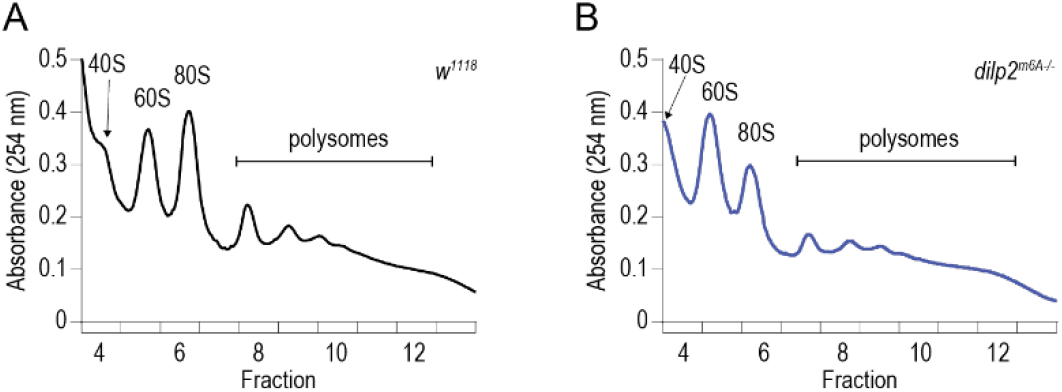
Polysome profiles of control and *dilp2^m6A−/−^* mutant flies. (A, B) Representative polysome profile from sucrose gradient of (A) control (w^1118^CS) and (B) *dilp2^m6A−/−^* mutant fly heads.

**Table S1.**
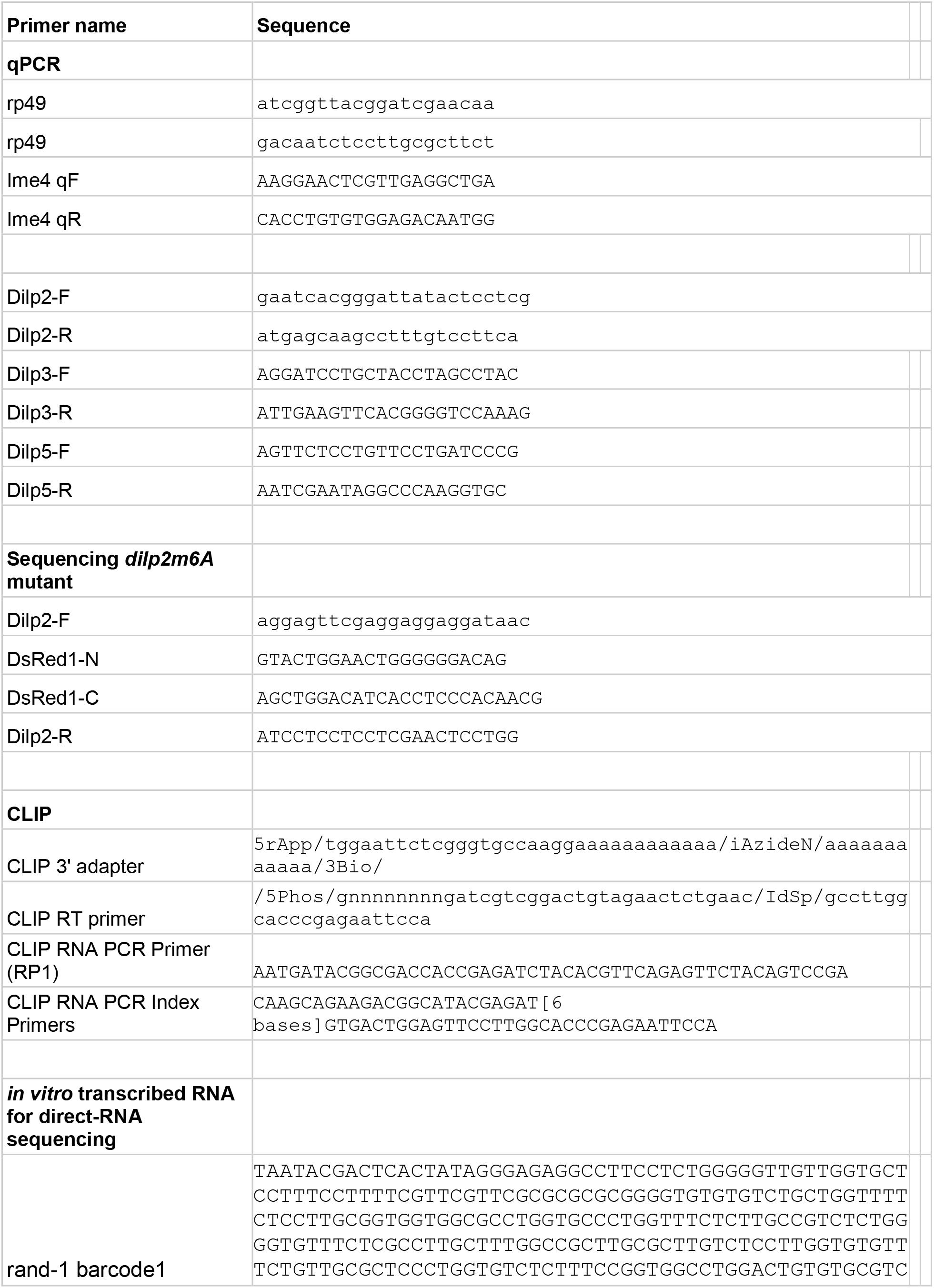
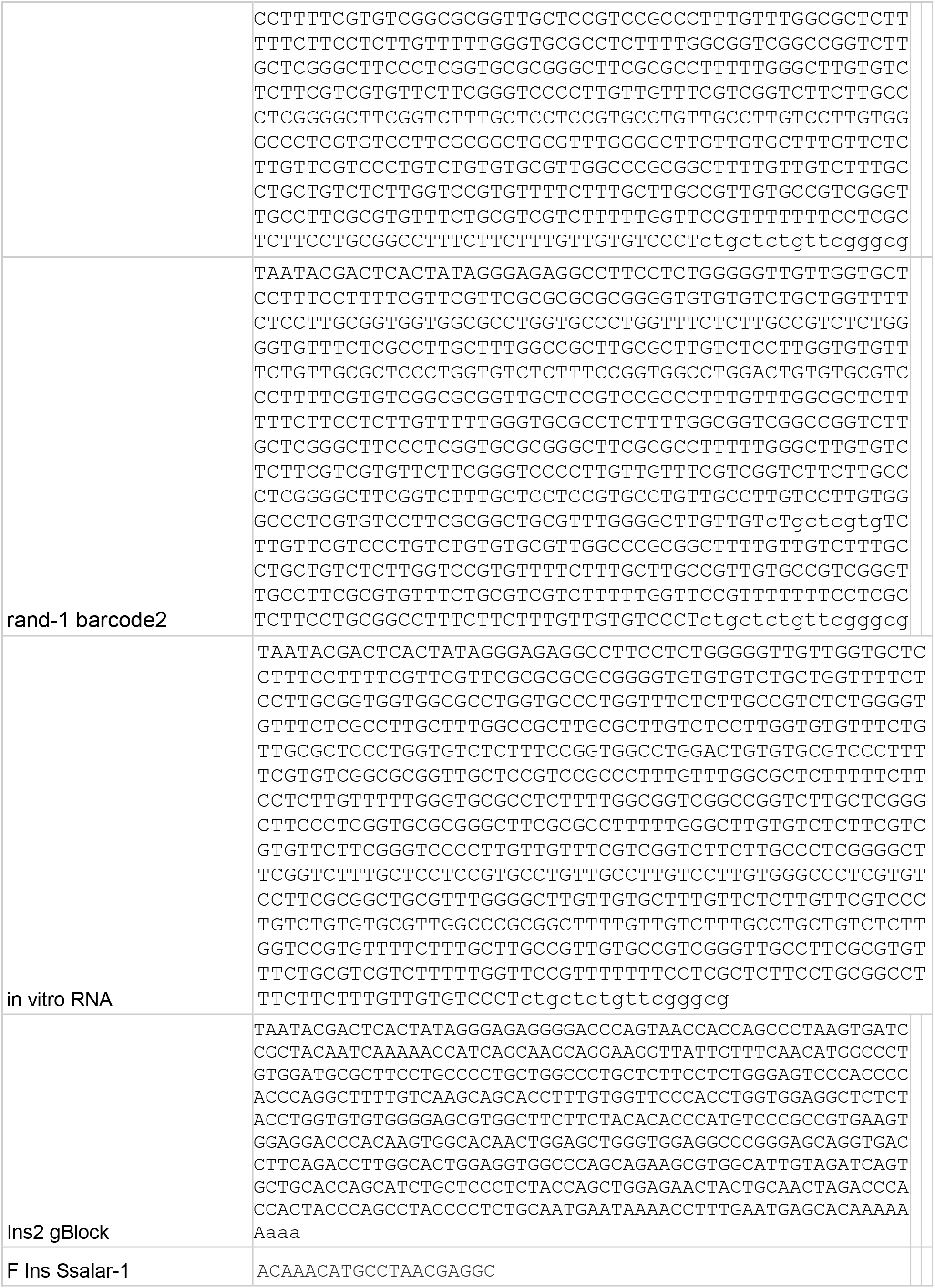
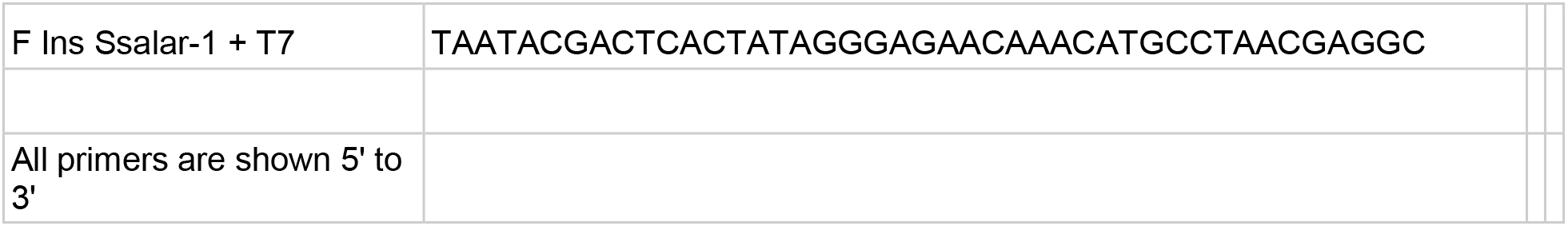
Oligonucleotide sequences. DNA sequences used for qPCR primers, Sanger sequencing, CLIP adaptors, and in vitro transcription of control RNAs.

**Table S2.**
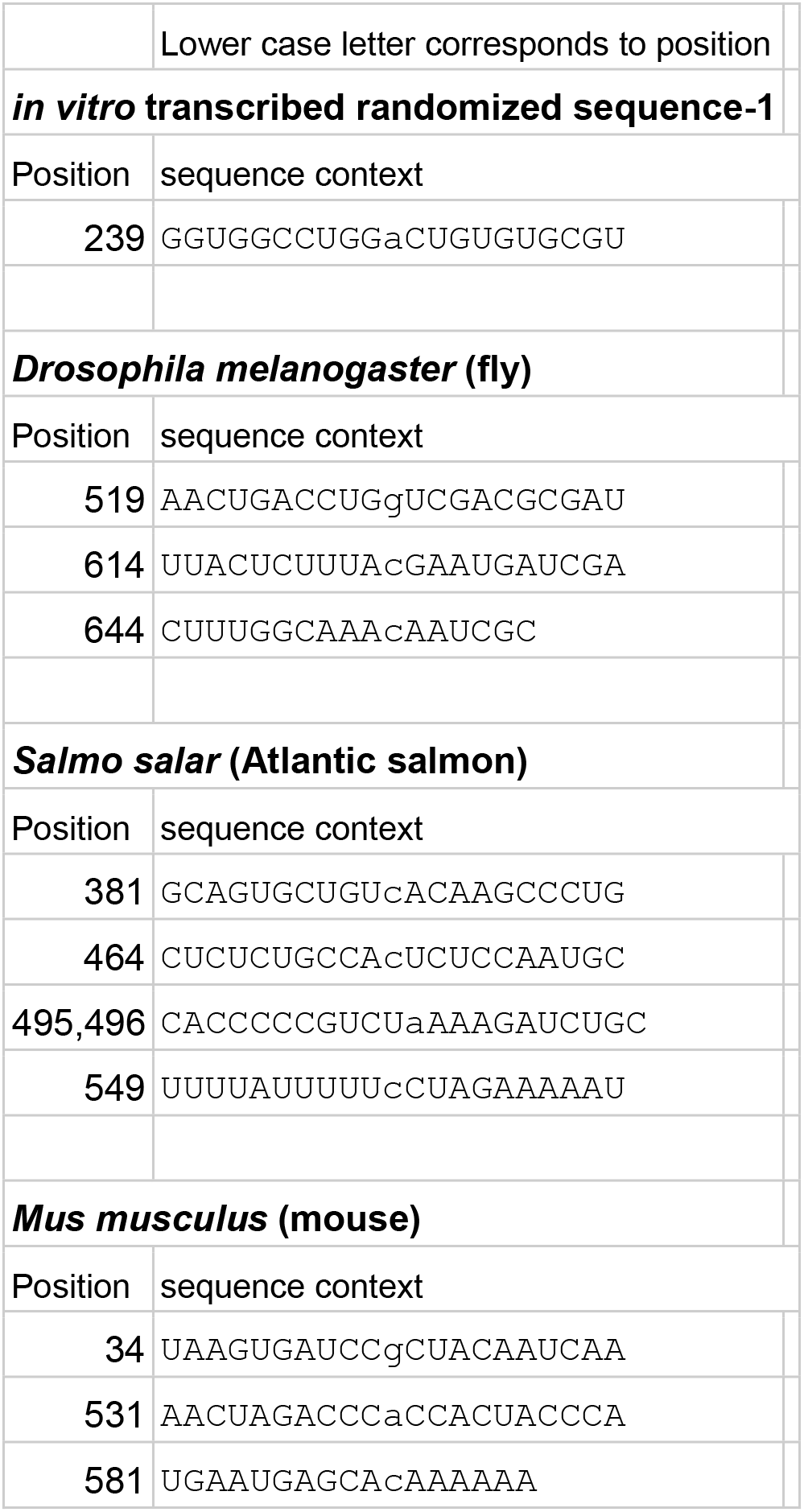
Sequence context of significant direct-RNA sequence differences. Sequences are listed for random *in vitro* transcribed RNA, and regions from native transcripts from each organism tested.

**Table S3.**
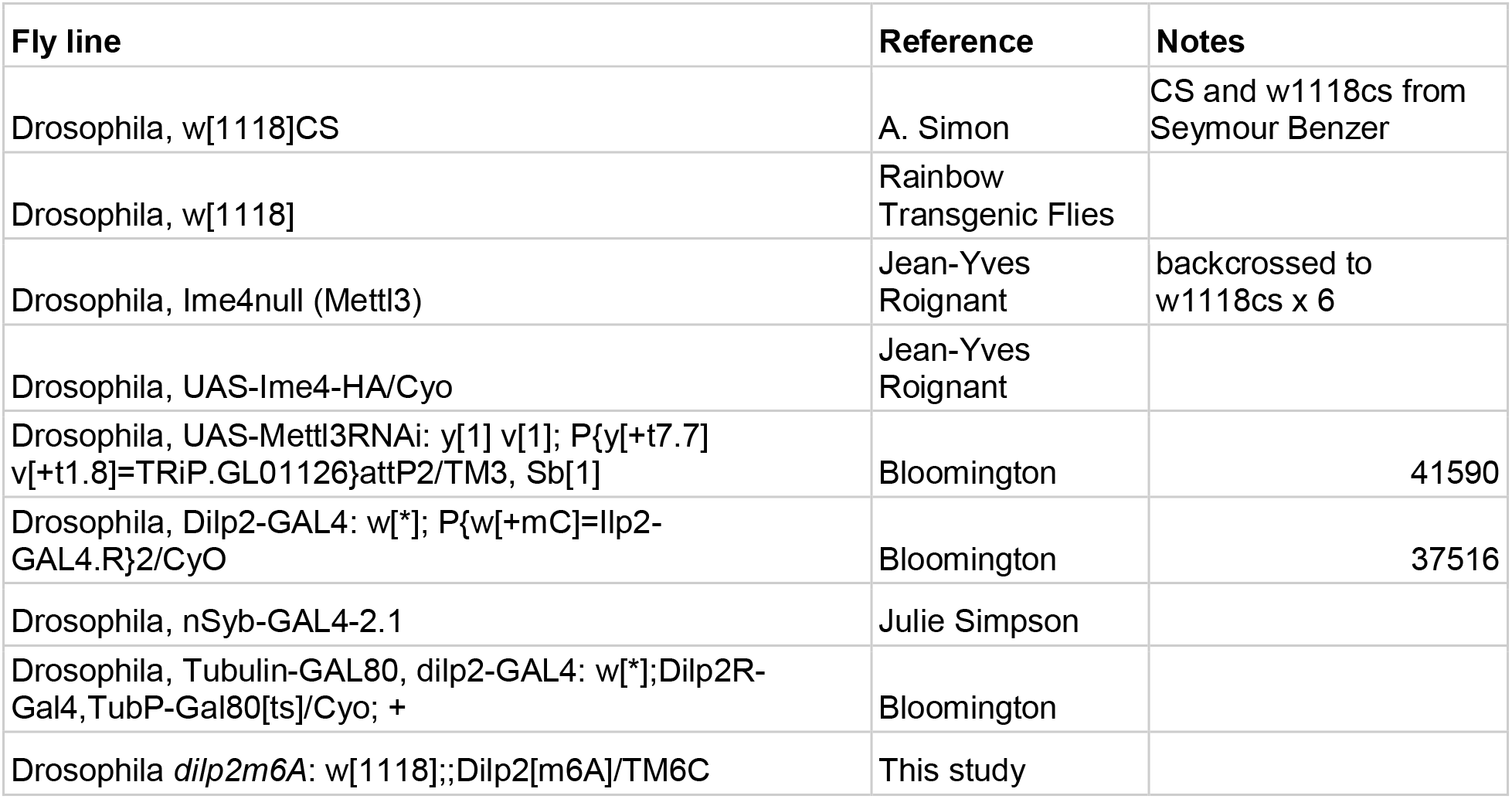
Fly stocks.

**Table S4. Summary of sequencing reads.**

**Data S1. List of CLIP peaks.**

Complete list of peaks from each biological replicate and the union of the data sets. CIMS output of C to T transitions.

